## Supplemental Material for "Simple Human Response Times are Governed by Dual Anticipatory Processes with Distinct and Distributed Neural Signatures"

**This PDF file includes:**

Figs. S1 to S3

Tables S1 to S2

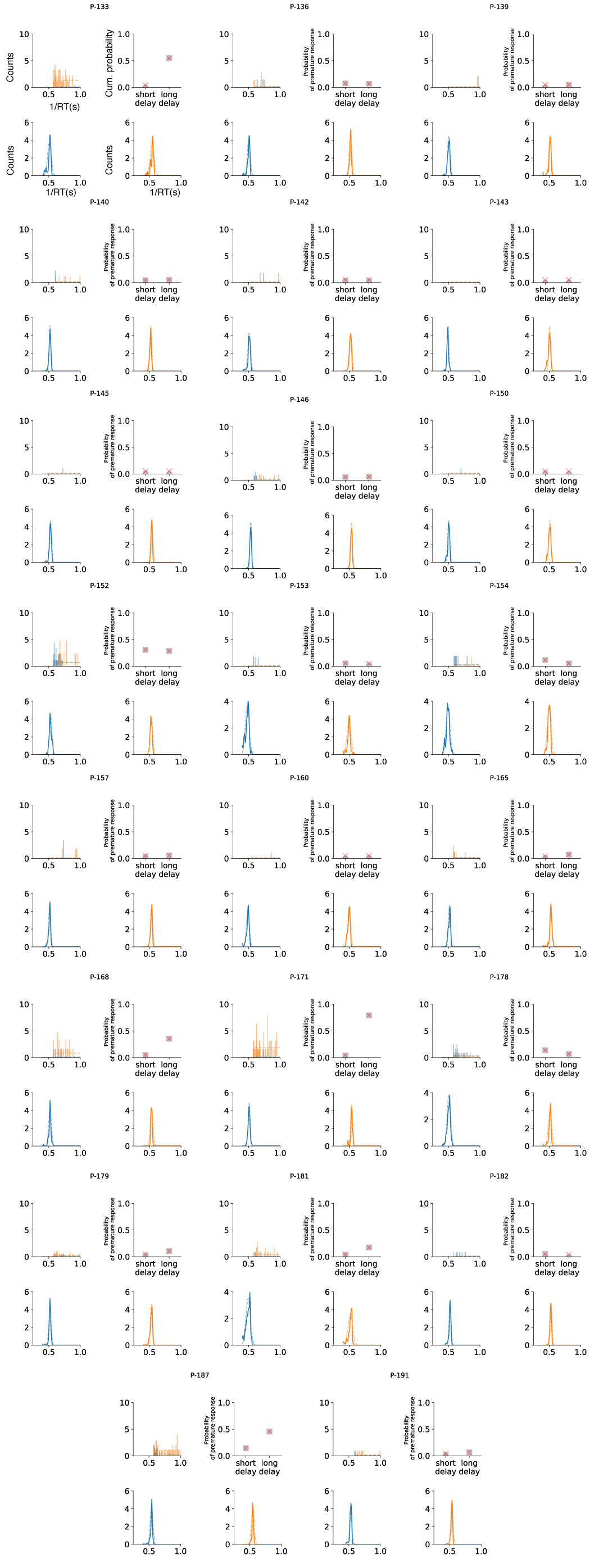

**Figure S1. Individual model fits** (**Participant ID 133–154**). Data from each participant are summarized in each block of four panels. The top-left panel shows the observed distribution of reciprocal RTs for premature false alarms during short-delay (blue vertical lines) and long-delay (orange vertical lines) trials, and the model-based uniform process underlying each distribution (dashed horizontal lines). The top-right panel shows the observed cumulative false-alarm rate for short- and long-delay trials (gray circles) and model predicted false-alarm rates (red crosses). Bottom panels show reciprocal RT distributions (solid line) and model fits (dashed line) on short-delay trials (blue, left panel) and long-delay trials (orange, right panel).

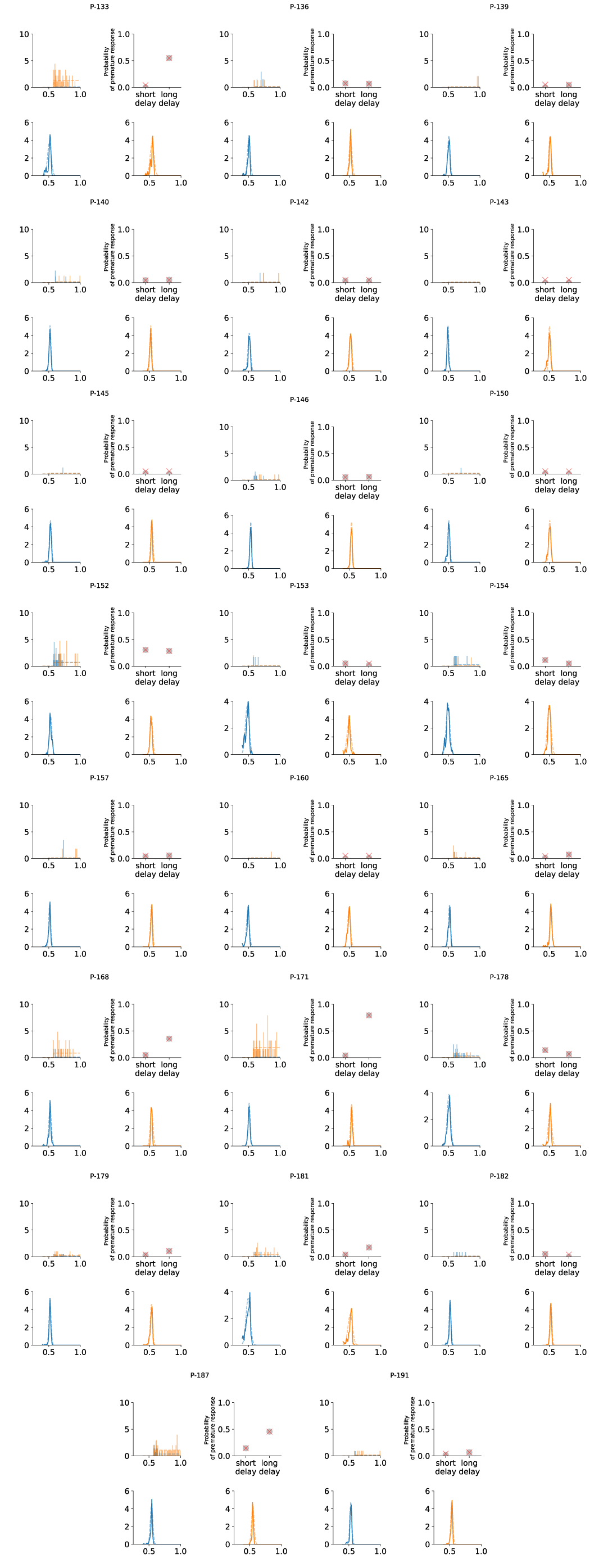

**Figure S1. Individual model fits** (**Participant ID 157-191**)

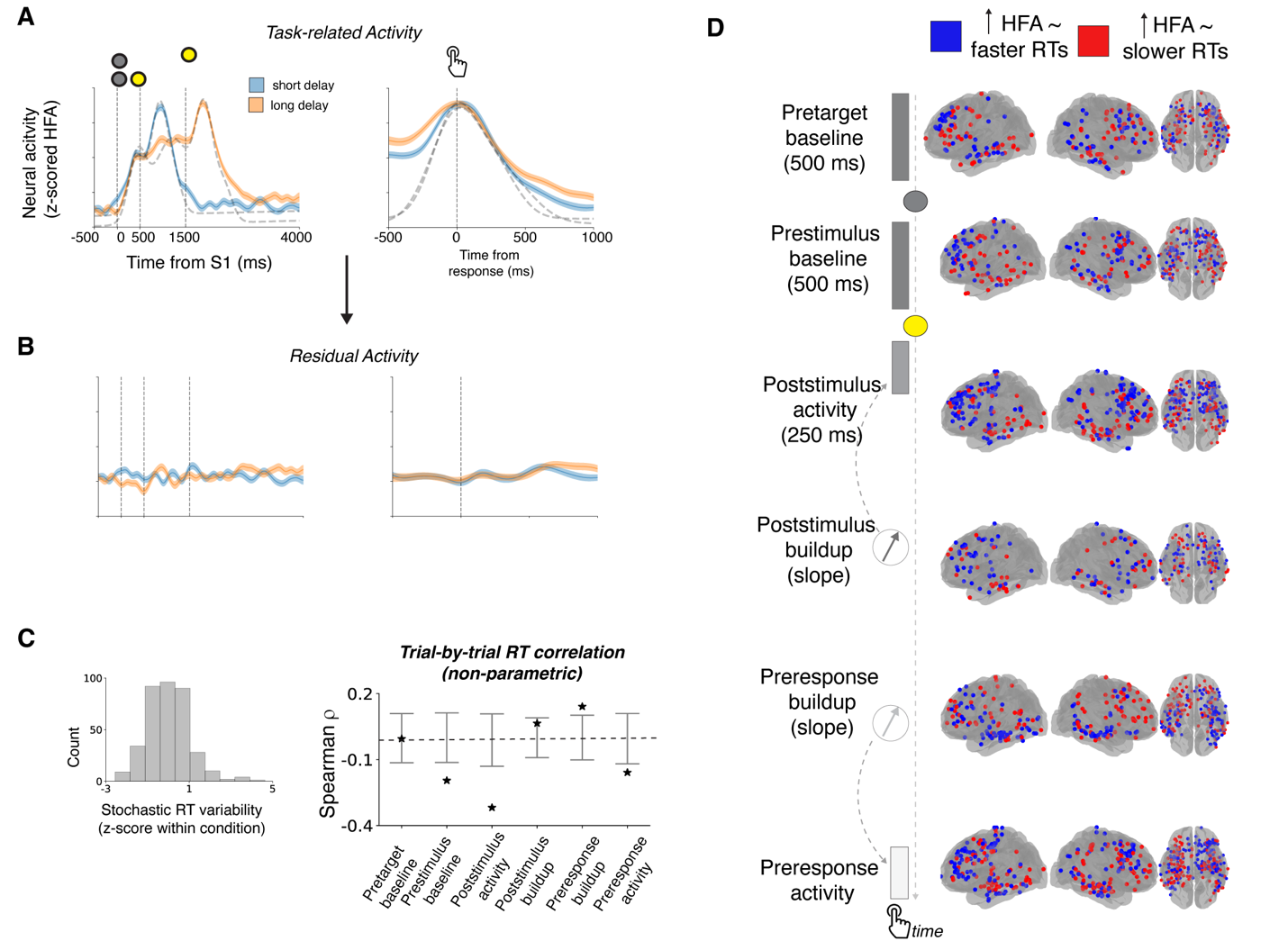

**Figure S2. Overview of methods used to relate each electrode’s activation function to endogenous RT variability**. (**A**) Example target-, stimulus-, and response-locked activation functions from a single electrode (same electrode as in Fig. 2C, without separation by RT). (**B**) Same as in A, but with stereotypical sensory and motor responses (determined via Gaussian fits to data across trials) removed. (**C**) Left panel shows the RT distribution measured in the same session as the neural data shown in **A** and **B**, with delay-related effects removed (z-scored separately per delay condition). Right panel shows how the residual activity shown in *B* measured at different times within a trial related to this RT variability. Asterisks indicate Spearman $\rho$*’s* computed from the data between trial-by-trial z-scored RT and neural activity, *p*<0.05; error bars indicate 95% confidence intervals. **(D)** Brain plots showing the anatomical distribution of RT-related neural representations (*p*<0.05, non-parametric tests). Blue electrodes indicate negative effects (relatively increased activity during fast RTs), red electrodes indicate positive effects (relatively increased activity during slow RTs). Each row depicts data from different task epochs, as indicated and as in **C**.

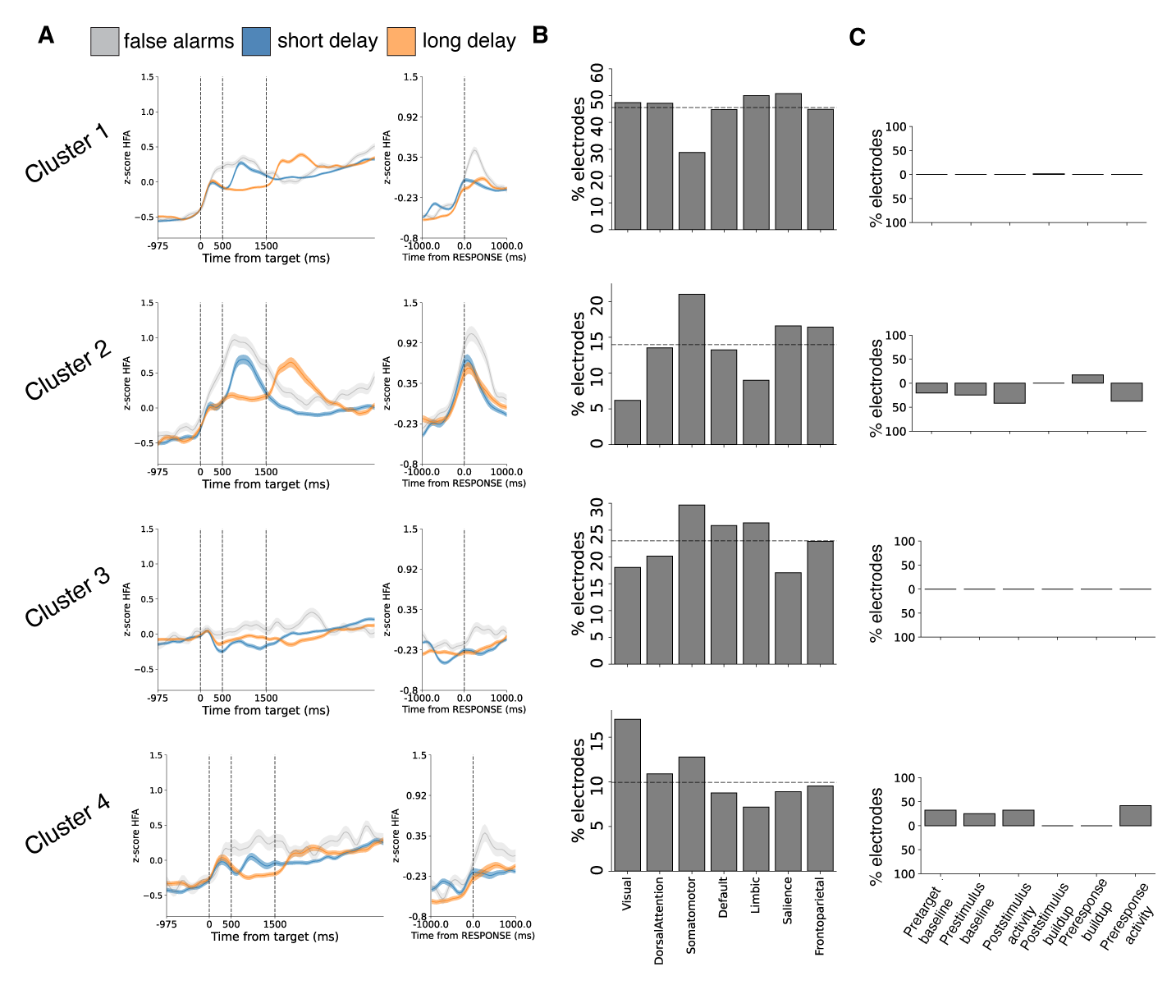

**Figure S3. Cluster descriptions**. Functional and anatomical properties of Clusters 1–4. (**A**) Average activation function (same format as Fig. 6B, but also including short-delay trials and response-locked activity). (**B**) Anatomical distribution of electrodes for each cluster across intrinsic brain networks. Height of bars indicate percentage of electrodes in each intrinsic brain network assigned to the given cluster. Horizontal dashed line is the expected percentage, assuming a uniform anatomical distribution across all networks. (**C**) Bar plot showing frequency of RT modulations for various task epochs (as illustrated in Figs. S2 and 4). Positive and negative values indicate frequency of electrodes showing positive effects and negative effects, respectively (as described in Fig. 3)

Table S1.

Participant characteristics

| Participant ID | Gender | Age at implant | Number of electrodes |
| --- | --- | --- | --- |
| 133 | Female | 52 | 51 |
| 136 | Female | 57 | 103 |
| 140 | Female | 48 | 39 |
| 142 | Male | 30 | 68 |
| 143 | Female | 30 | 97 |
| 145 | Male | 21 | 96 |
| 146 | Male | 19 | 96 |
| 150 | Male | 16 | 90 |
| 152 | Female | 33 | 79 |
| 153 | Female | 56 | 92 |
| 154 | Female | 43 | 199 |
| 157 | Male | 25 | 145 |
| 160 | Female | 46 | 148 |
| 165 | Female | 20 | 93 |
| 168 | Male | 27 | 170 |
| 171 | Male | 49 | 160 |
| 178 | Female | 38 | 140 |
| 179 | Female | 19 | 140 |
| 181 | Female | 31 | 138 |
| 182 | Female | 25 | 118 |
| 184 | Male | 22 | 161 |
| 187 | Male | 24 | 74 |
| 191 | Female | 32 | 112 |

**Table S2.**

Participant Behavior

| Participant | n. trials | n. sessions | mean RT on long delay (ms) | std RT on long delay (ms) | mean RT on short delay (ms) | std RT on short delay (ms) | false alarm rate on short delay | false alarm rate on long delay | lapse rate on short delay | lapse rate on long delay |
| --- | --- | --- | --- | --- | --- | --- | --- | --- | --- | --- |
| 133 | 248 | 2 | 352.32 | 100.83 | 465.22 | 122.59 | 0.00 | 0.28 | 0.03 | 0.01 |
| 136 | 236 | 1 | 425.99 | 53.87 | 489.51 | 95.57 | 0.06 | 0.08 | 0.00 | 0.03 |
| 139 | 119 | 1 | 455.97 | 103.75 | 475.06 | 76.10 | 0.00 | 0.07 | 0.05 | 0.07 |
| 140 | 177 | 1 | 428.92 | 52.22 | 422.20 | 53.46 | 0.01 | 0.15 | 0.00 | 0.00 |
| 142 | 115 | 1 | 425.36 | 59.68 | 460.05 | 87.90 | 0.04 | 0.03 | 0.02 | 0.03 |
| 143 | 110 | 1 | 487.45 | 92.09 | 531.96 | 56.92 | 0.00 | 0.02 | 0.02 | 0.00 |
| 145 | 167 | 1 | 364.29 | 31.22 | 396.15 | 64.73 | 0.01 | 0.05 | 0.01 | 0.00 |
| 146 | 234 | 1 | 375.09 | 50.71 | 373.64 | 46.95 | 0.01 | 0.16 | 0.01 | 0.00 |
| 150 | 167 | 1 | 467.87 | 76.21 | 479.84 | 68.42 | 0.01 | 0.05 | 0.00 | 0.00 |
| 152 | 164 | 1 | 398.42 | 53.55 | 389.61 | 72.92 | 0.08 | 0.57 | 0.01 | 0.00 |
| 153 | 124 | 1 | 529.43 | 123.42 | 595.26 | 141.85 | 0.00 | 0.12 | 0.11 | 0.05 |
| 154 | 120 | 1 | 501.74 | 90.12 | 533.73 | 122.79 | 0.02 | 0.10 | 0.02 | 0.01 |
| 157 | 116 | 1 | 379.88 | 46.81 | 444.45 | 65.56 | 0.03 | 0.09 | 0.00 | 0.02 |
| 160 | 174 | 1 | 523.43 | 81.61 | 547.32 | 103.38 | 0.00 | 0.05 | 0.09 | 0.01 |
| 165 | 166 | 1 | 400.61 | 95.09 | 435.12 | 63.88 | 0.00 | 0.05 | 0.00 | 0.00 |
| 168 | 130 | 1 | 373.69 | 28.14 | 422.86 | 82.53 | 0.00 | 0.28 | 0.00 | 0.00 |
| 171 | 157 | 1 | 366.53 | 72.75 | 464.91 | 62.67 | 0.02 | 0.59 | 0.01 | 0.00 |
| 178 | 260 | 2 | 451.93 | 91.35 | 478.19 | 110.55 | 0.02 | 0.20 | 0.05 | 0.05 |
| 179 | 403 | 3 | 390.27 | 79.01 | 423.30 | 72.34 | 0.02 | 0.09 | 0.01 | 0.01 |
| 181 | 263 | 2 | 451.22 | 139.21 | 539.17 | 152.55 | 0.02 | 0.17 | 0.11 | 0.05 |
| 182 | 225 | 2 | 399.13 | 43.01 | 402.90 | 61.81 | 0.03 | 0.05 | 0.00 | 0.01 |
| 187 | 290 | 2 | 323.76 | 40.88 | 354.84 | 77.84 | 0.00 | 0.49 | 0.00 | 0.01 |
| 191 | 231 | 2 | 372.18 | 50.44 | 386.12 | 63.19 | 0.00 | 0.12 | 0.00 | 0.00 |
